# Independent component analysis provides clinically relevant insights into the biology of melanoma patients

**DOI:** 10.1101/395145

**Authors:** Petr V. Nazarov, Anke K. Wienecke-Baldacchino, Andrei Zinovyev, Urszula Czerwińska, Arnaud Muller, Dorothée Nashan, Gunnar Dittmar, Francisco Azuaje, Stephanie Kreis

## Abstract

The integration of publicly available and new patient-derived transcriptomic datasets is not straightforward and requires specialized approaches to deal with heterogeneity at technical and biological levels. Here we present a methodology that can overcome technical biases, predict clinically relevant outcomes and identify tumour-related biological processes in patients using previously collected large reference datasets. The approach is based on independent component analysis (ICA) – an unsupervised method of signal deconvolution. We developed parallel consensus ICA that robustly decomposes merged new and reference datasets into signals with minimal mutual dependency. By applying the method to a small cohort of primary melanoma and control samples combined with a large public melanoma dataset, we demonstrate that our method distinguishes cell-type specific signals from technical biases and allows to predict clinically relevant patient characteristics. Cancer subtypes, patient survival and activity of key tumour-related processes such as immune response, angiogenesis and cell proliferation were characterized. Additionally, through integration of transcriptomes and miRNomes, the method identified biological functions of miRNAs, which would otherwise not be possible.

## INTRODUCTION

Genomic and transcriptomic research has accumulated a vast collection of publicly available cancer-related data. Integrating this information still poses a considerable obstacle as genomic and transcriptomic data from cancer patients are characterized by significant heterogeneity at two levels. First, results are generally collected using different sample preparation protocols and transcriptome analysis platforms and are then interrogated by constantly changing techniques. Although these techniques have improved on accuracy, sensitivity or genome coverage, they restrain backward compatibility, e.g., expression level analysis has evolved from qPCR through microarrays toward NGS sequencing in the last 10-15 years. Second, collected patient samples are intrinsically heterogeneous at tissue and cellular levels. Bulk analysis of transcriptomes can mask different types of heterogeneity in the sample as tumour biopsies contain many cell types that are mixed in different proportions. Furthermore, there are well-documented variations of tumour cells within the same neoplasia, which can conceal low abundant, but critical cell subtypes such as drug-resistant tumour cells. These facts plus the lack of standardised analysis pipelines limits discoveries and can lead to erroneous clinical conclusions (Patel, Tirosh et al., 2014, Zhao, Hemann et al., 2014). The experimental approach to resolve the complex issue of working with heterogeneous cancer samples involves physical separation of tissue into homogeneous cell populations or even single cells (by cell sorting, single cell technologies or microdissection) before the actual measurement (Legres, Janin et al., 2014, Patel et al., 2014, Weaver, Tseng et al., 2014). Technologically, this is an expensive and laborious task, which is not yet accessible routinely and which can introduce experimental errors (Debey, Schoenbeck et al., 2004, Shannon, Balshaw et al., 2014).

Alternatively, computational approaches can be applied to separate or deconvolute multivariate signals from different cell types, accounting for variable biopsy sample composition and intra-tumour heterogeneity. One of the most promising methods of assumption-free transcriptome deconvolution is independent component analysis (ICA). This method originated from the domain of signal processing aiming at detecting individual components from a complex mix of mutually independent non-Gaussian signals. It allows to identify sources of transcriptional signals, cluster genes into functional groups and cell type-related signatures (Biton, Bernard-Pierrot et al., 2014, Teschendorff, Journee et al., 2007) and deduce interactions between biological processes (Lee & Batzoglou, 2003). Importantly, ICA results are concordant with clinical data, such as cancer subtypes (Biton et al., 2014) and survival, suggesting ICA as a potentially useful tool for patient diagnostics and prognostics. The method can also recognise and remove biologically irrelevant signals such as technical factors and/or confounders. Therefore, such an approach can make use of large sets of high-throughput transcriptomic (or other) data that were collected through different stages of technological progress.

For cancer research, one of the most valuable data sources is The Cancer Genome Atlas (TCGA), which holds over 10 000 patient-derived samples including various levels of omics data: DNA, RNA, and proteins. The data have been continuously collected using massive financial and scientific efforts. Now, the question arises if this resource can also be used to support clinicians in making rapid and accurate assessments leading to tailored treatments for individual cancer patients. Therefore, methods that can incorporate newly generated ‘omics’ data (further addressed as “investigation dataset”) in large public reference datasets are required for improved diagnostics of new patients. Here we propose an ICA-based method to combine reference and investigation datasets for patient diagnostics and detailed inspection of biological processes in cutaneous melanoma (Hayward, Wilmott et al., 2017).

Melanoma arises through the malignant transformation of melanocytes and presents a very aggressive form of skin cancer with increasing global case numbers (https://www.cancer.gov/aboutcancer/understanding/statistics). Approximately 50-60% of patients express mutated BRAF or NRAS, rendering the BRAF kinase signalling cascade constitutively active, impacting on both MAPK and PI3K/AKT pathways (Holderfield, Deuker et al., 2014). The introduction of specific kinase inhibitors of the activating BRAF kinase mutation and downstream MEK kinases, and the further use of immune checkpoint inhibitors have improved patients’ overall survival (Luke, Flaherty et al., 2017). However, resistance or severe side effects can arise rapidly against both types of therapies, leaving hardly any treatment options for the majority of advanced stage melanoma patients resulting in a low 5-year survival rate (Schadendorf, Fisher et al., 2015, Sullivan & Flaherty, 2013). Melanoma’s extremely high mutation rate (>10 somatic mutations/Mb) and the concomitant genetic heterogeneity make it difficult to distinguish true cancer driver genes from noise and low frequency mutations in bulk samples using current technologies (Lawrence, Stojanov et al., 2013, Zhang, Dutton-Regester et al., 2016). Nevertheless, according to mutational hotspots, melanoma patients have recently been allocated to four groups: BRAF, NF1, RAS or Triple WT (wild type), while the analysis of gene expression data resulted in three functional groups: “immune subclass”, “keratin subclass” and “MITF-low subclass”, which according to the authors have implications for patient survival (Cancer Genome Atlas, 2015). Interestingly, the majority of primary melanomas belonged to the “keratin subclass” having a worse prognosis than the other two subclasses.

In this study, we used the SKCM TCGA cohort with over 470 patients diagnosed with cutaneous melanoma as the reference dataset. Two layers of sequencing data were considered and integrated: mRNA and microRNA (miRNA). The investigation dataset included a small cohort of three primary melanoma tumours and two controls: matched cancer patient-derived normal skin and NHEM cells (normal human epidermal melanocytes). First, for the reference cohort, we demonstrated that ICA deconvolution was successfully applied to classify patients based on their tumour subtypes and to build the hazard score that predicts patient survival. The hazard score was then tested using an independent validation cohort of 44 patients, obtained by microarray gene expression technology. The strong difference between reference RNA-seq data and microarray-derived validation datasets was overcome by our method. Next, the investigated dataset was studied in depth and key processes involved in cancer aetiology were detected: immune response and inflammation, angiogenesis, self-sufficient cell proliferation and other.

We show here that consensus ICA can integrate data from different biological sources, time points and platforms to predict clinically important characteristics of cancer in a bias-free, unsupervised and potentially automatable fashion, suggesting consensus ICA as a useful module of future clinical “one-click” support systems.

## RESULTS

### ICA of combined data sets can remedy technical biases

#### Reference, validation and investigation datasets

Using publicly available data sets for diagnostic/prognostic purposes on new patients is often hampered by the fact that different technical platforms, data formats and analysis pipelines are used to generate the reference data and the patient samples to be investigated. Thus, reference and newly generated data do not come from the same distribution, which often introduces strong technical biases despite different normalisation schemes. It has recently been shown that ICA, when applied to heterogeneous datasets affected by technical factors, can identify such biases and single out one or several components to account for them (Biton et al., 2014, Taroni & Greene, 2017). As soon as the technical bias is captured by an independent component, it is isolated from the core data, which can then be analysed in a more accurate way.

In this study, graphically outlined in Figure 1, we used public TCGA data as the reference dataset, published microarray data (Bogunovic, O’Neill et al., 2009) as a validation set and an investigation set based on clinical samples – three primary melanoma tumours (denoted P2PM, P4PM, P6PM) and two control samples (matched normal skin P4NS and NHEM cell line) as described in Table 1. We developed a method to assign a prognostic hazard score based on the reference set and validated our approach on an independent public microarray data set, followed by testing the method on clinical samples. The underlying ICA was done on single or combined datasets (reference only, reference/validation and reference/investigation) by the developed consensus ICA method.

**Table 1.**
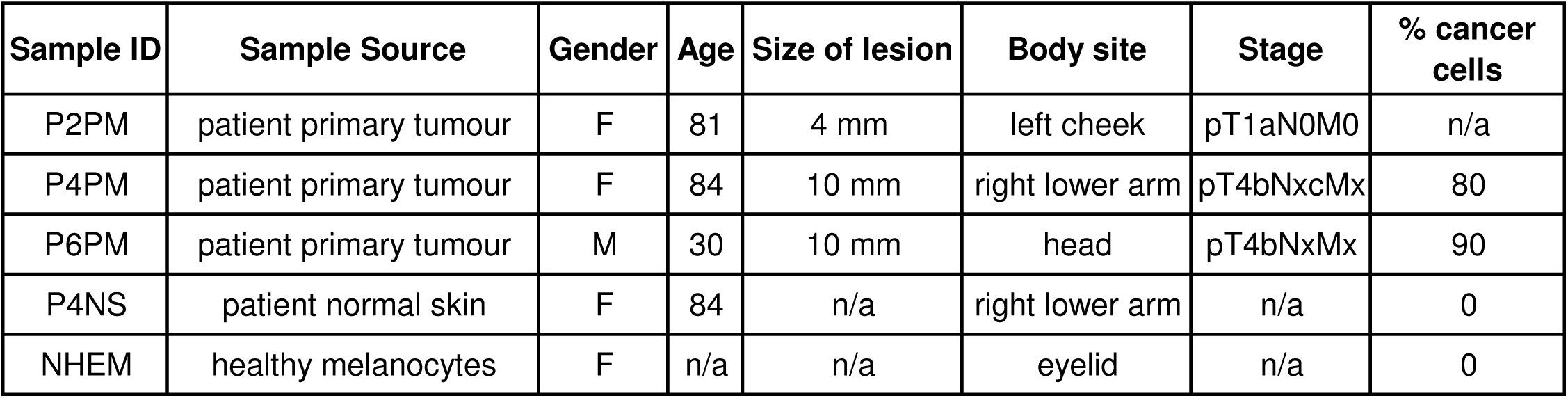
Parameters of clinical samples and controls in the investigation dataset.

**Figure 1.**
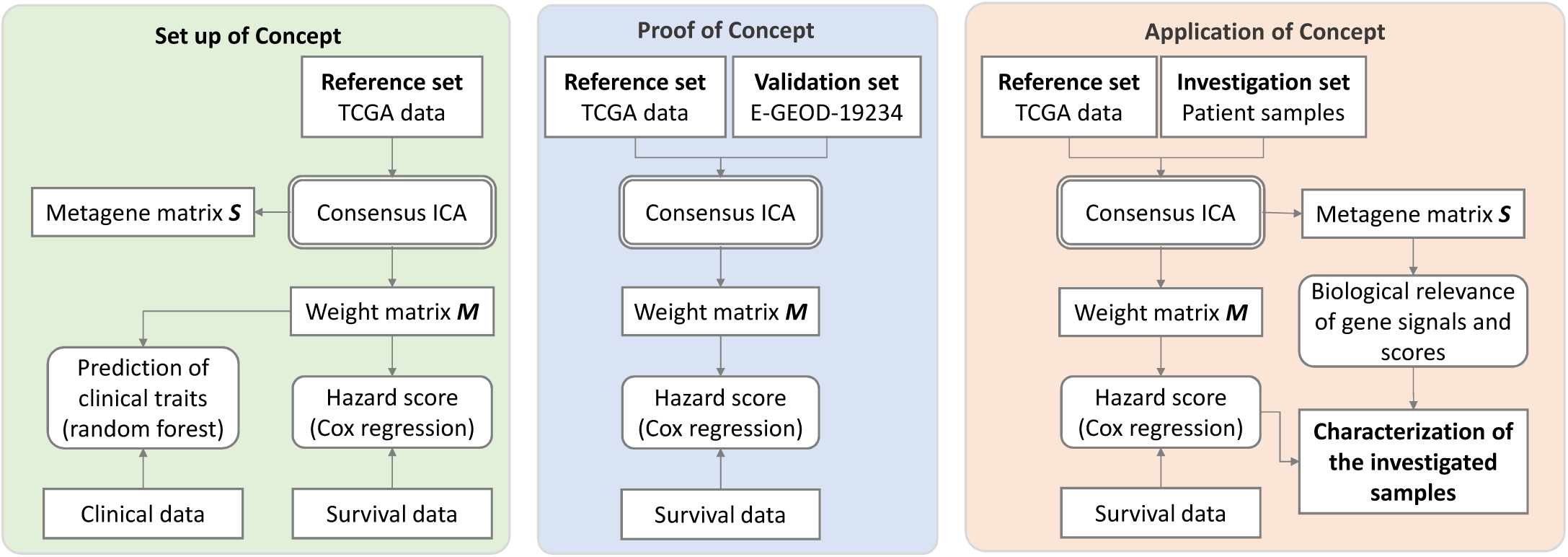
Schematic workflow of ICA application to the reference, validation and investigation datasets. Left panel (green): preliminary ICA of reference TCGA data. We established the technique, investigated the RNA-seq measures (counts and FPKM), selected the number of components, showed that weight matrix ***M*** can be used for patient classification and set up a hazard score. Middle panel (blue): the developed hazard score was tested on the additional validation dataset. Left panel (red): application of the method on an unpublished investigation dataset of 5 samples: 3 primary tumours, one normal skin and one NHEM cell line. Transcriptome and miRNome data integration and in-depth investigation of the biologically relevant signals seen in ***S***-matrix were performed.

#### Stability of deconvolution

ICA was applied to two types of transcriptomic data: mRNA and miRNA expression. Based on the reasoning provided in Supplementary Methods, 80 independent components were used for the deconvolution of mRNA data (named RIC1-80) and 40 for miRNA data (denoted as MIC1-40). ICA was run 1000 times in order to achieve robust results. Of 80 RICs, 26 showed high reproducibility (mean coefficient of determination between the detected metagenes R^2^ > 0.8 after 1000 runs of ICA) while 23 had reasonable reproducibility (0.5 < R^2^ ≤ 0.8). Among 40 MICs, 23 had R^2^ > 0.8 and 13 – 0.5 < R^2^ ≤ 0.8. As mentioned in Supplementary Methods, RICs were automatically reoriented in order to make the biologically relevant sets of genes contribute positively to the matrix of metagenes ***S***. MICs were reoriented based on the negative sign of correlation with RICs.

#### ICA identifies technical biases

The combined reference/investigation dataset profiled by RNA-seq is presented in the space of two first principal components (Figure 2A) and of two selected independent components (Figure 2B). The two principal components included 33% of total variability and mainly reflected technical effects: PC1 was linked to the RNA-seq library size (data not shown) and PC2 segregates reference and investigation data. Among all RICs, the components that reflected data clustering according to gender (RIC3) and sample type (primary or metastatic, RIC5) were chosen. Both components were detected with high reproducibility (R^2^_RIC3_ = 0.996, R^2^_RIC5_ =0.993). The investigation data were clearly integrated within the reference data and showed reasonable clustering in Figure 2B. P2PM and P4PM samples originated from primary tumours of female patients, P4NS normal skin corresponding to P4PM samples and NHEM was extracted from eyelid skin of a female. P6PM was a primary tumour of a male patient. Functional analysis showed that genes involved in RIC5 participate in keratinocyte-specific functions and thus weights of RIC5 could be used as a marker of keratinocyte presence. Indeed, the vast majority of metastatic samples had low values of RIC5 weights, while primary tumours showed high values. NHEM (pure melanocytes) are devoid of keratinocytes and therefore clustered with metastatic tissues.

**Figure 2.**
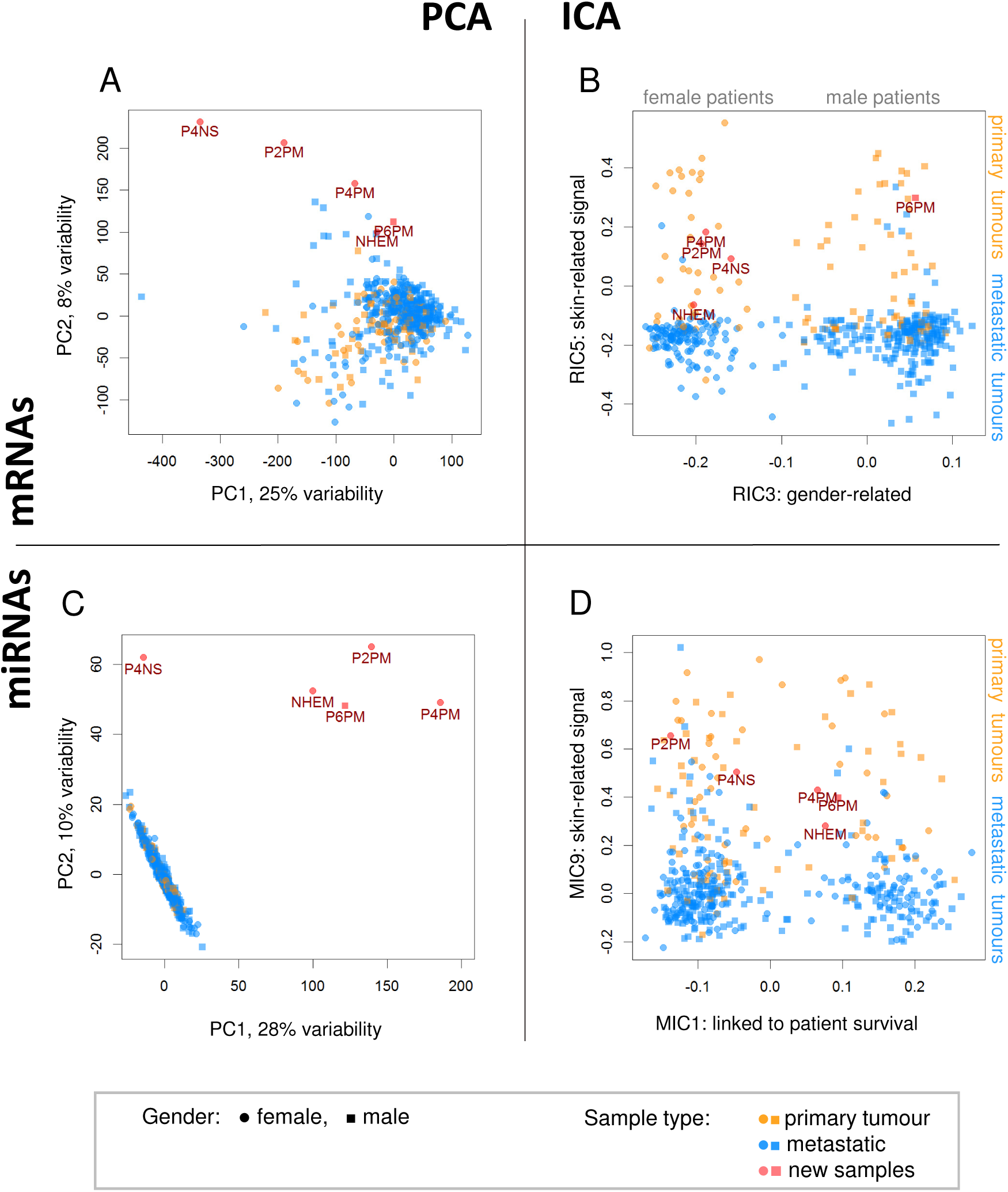
Data variability captured by the first components of PCA (A) and two selected components of ICA (B) in gene expression data. Independent components were selected based on the predictive power of their weights for patient gender (RIC3) and sample type (RIC5). MiRNA data showed even higher discrepancy comparing miRNA-seq and qPCR results by PCA (C). However, in the space of independent components (MIC1 and MIC9), the samples studied by miRNA-seq and qPCR overlap (D).

An even stronger normalisation effect of ICA was observed for miRNA data. As mentioned above, the techniques used for miRNA detection and quantification were different: miRNA-seq for the reference dataset and whole miRNome qPCR arrays for the new samples. PCA showed strong differences between log_2_ transformed counts and inverted Ct values (Figure 2C). However, in the space of independent components, the new samples were properly located again (Figure 2D). Here, two miRNA components MIC1 and MIC9 were depicted. MIC1 showed a strong relation to survival (Cox-based log rank p-value=9.4e-4) while MIC9 was correlated with the skin-related signal of RIC5.

### ICA yields clinically relevant information

#### ICA as a feature-selection method for sample classification

As observed for patient gender and sample type in Figure 2B, the weights of the components can be used as features with predictive potential. We investigated whether clinical factors could be predicted by ICA deconvolution, only analysing RICs for simplification. Three factors were selected for the report: gender, sample type and RNA cluster, that could be considered as cancer subtype and was previously introduced in (Cancer Genome Atlas, 2015). We validated the random forest classification directly on the reference set using leave-one-out cross-validation (LOOCV), as described in Material and Methods. Then, the new samples were classified and the results are presented in Table 2. Gender and sample types were accurately predicted for all the new samples but NHEM cells were considered metastatic: the best location predictors were weights linked to the transcriptional signal of keratinocytes, which was low in metastatic tumours and also in this primary cell line.

**Table 2.**
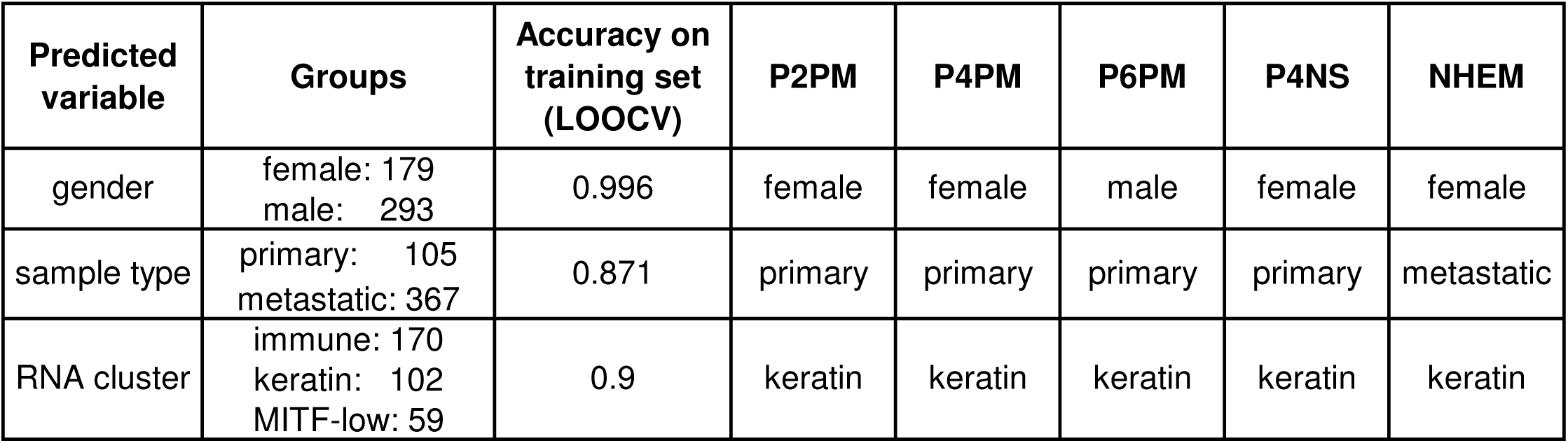
Performance of ICA-based feature extraction. Accuracy was calculated using leave-one-out cross-validation on the TCGA reference set; predictions for the investigation set are reported.

| Predicted variable | Groups | Accuracy on training set (LOOCV) | P2PM | P4PM | P6PM | P4NS | NHEM |
| --- | --- | --- | --- | --- | --- | --- | --- |
| gender | female: 179<br>male: 293 | 0.996 | female | female | male | female | female |
| sample type | primary: 105<br>metastatic: 367 | 0.871 | primary | primary | primary | primary | metastatic |
| RNA cluster | immune: 170<br>keratin: 102<br>MITF-low: 59 | 0.9 | keratin | keratin | keratin | keratin | keratin |

#### ICA provides prognostic features linked to patient survival

Next, prognostic abilities of the ICA weights were examined by a Cox regression model. All components, their significance and log hazard ratios (LHRs) are summarised in Supplementary Tables S1 and S2. Eleven RICs and 3 MICs were found significantly linked to patient survival after multiple testing adjustment (FDR-adjusted log rank p-value for Cox regression < 0.05). Among them, 6 RICs and 2 MICs showed very high stability of R^2^>0.8: RIC2, RIC4, RIC5, RIC7, RIC25, RIC75, MIC1 and MIC20. Although these 11 RICs and 3 MICs were statistically linked to survival in our reference set, the predictive power of any of them may have not been sufficient to predict survival of new patients. Therefore, we combined the weights of these components into a hazard score (HS) as described in Material and Methods. Combined HS showed high significance (Cox log rank p-value = 2.2e-13) for the TCGA dataset.

In order to validate the proposed hazard scoring approach on an independent cohort of patients, we repeated the ICA on the combined TCGA RNA-seq data (reference set) and microarray data E-GEOD-19234 (additional validation set). Notably, the metagenes of the new decomposition were strongly correlated with the metagenes from the initial reference-set; only ICA and 44 of them showed R^2^>0.5 (95% prediction interval for R^2^ values between metagenes of the same decomposition was: 0<R^2^<0.004). The components that showed a significant (adj.p-value<0.05) link to survival on the reference set were then used to compose HS for the validation data and also showed significant prognostic properties (log rank p-value of 0.0013); Kaplan-Meyer plot shown in Figure 3. The developed HS separated patients with low hazard (only one death among 7 patients, blue line in validation set, Figure 3) from the group of patients with a high hazard score.

**Figure 3.**
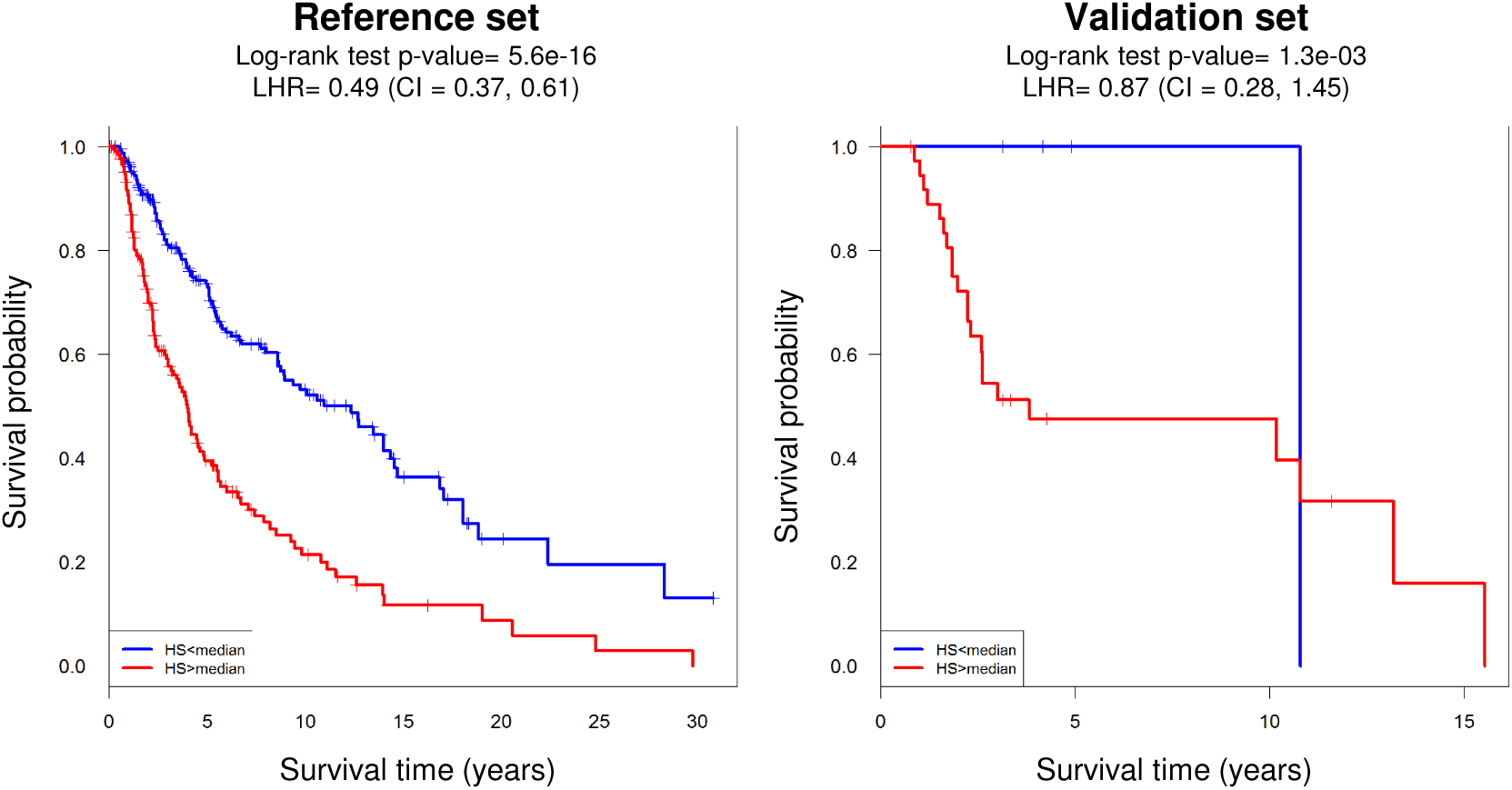
ICA-based hazard score (HS) can predict patient survival. Performance of the score on the TCGA reference set (A). Validation of the hazard score on the independent cohort composed of 44 metastatic melanoma patients (B). Cox regression log hazard ratio (LHR), together with its 95% C.I. and log rank p-value, are reported. In order to visualize the results as Kaplan-Meyer curves, patients were divided into two groups by their HS (low - blue and high - red).

For the three primary melanoma samples from the investigation set, the calculated HS was the lowest for the P2PM sample (HS=1.16) and the highest for P6PM (HS=1.92). This was in agreement with clinical observations, as patient P6 suffered from a very aggressive form of melanoma and deceased shortly after sample collection. From the quantitative results obtained from the validation dataset and qualitative differences observed for the investigation dataset, we concluded that weights of independent components can be combined into a hazard score, suitable to predict patient survival.

### Independent components provide information about biological processes in tumours

#### General strategy

The most challenging part of ICA is assigning components to specific biological processes, cell types and technical factors. The approach we have taken is outlined in Supplementary Figure S1 and the automatically generated reports describing the components can be found in the Supplementary Results. Briefly, the weights of components were linked to clinical factors using ANOVA and to survival by Cox regression. RIC metagenes were submitted to an over-representation analysis, resulting in enriched categories (GO terms (Alexa & Rahnenfuhrer, 2016), cell types, chromosome locations, etc. (Kuleshov, Jones et al., 2016)). Next, we compared the detected RIC metagenes to previously published ones derived using ICA from bladder cancer transcriptomes (Biton et al., 2014) and to the *LM22* leukocyte gene signature matrix (Newman, Liu et al., 2015). Comparison to the published metagenes showed common signals between melanoma and bladder cancers (Supplementary Figure S2A), while the LM22 signature matrix helped to better identify types of leukocytes (Supplementary Figure S2B). LM22 contains 547 genes and distinguishes 22 human hematopoietic cell types, including several T cells, B cells, NK cells and others. The biological origin of the detected signals was confirmed using melanoma single-cell data reported by Tirosh et al. (Tirosh, Izar et al., 2016) – correlation of several components is shown in Supplementary Figure S3. MICs were linked to RICs using correlation between weights of the components and literature mining. Genomic locations of MICs were further used to find cytogenic bands enriched with top-contributing miRNAs. Finally, we used the weights of the components linked to biologically relevant signals in order to acquire new knowledge about processes in the investigated samples.

#### Immune components

The biggest cluster of RICs was linked to immune cells and immune response. Based on functional annotation it included seven components: RIC2, RIC25, RIC27, RIC28, RIC37, RIC57 and MIC20. RIC2, RIC25 and RIC27 showed correlated weight profiles between themselves and with RIC74, RIC79 and MIC20 (Figure 4A, Supplementary Results).

**Figure 4.**
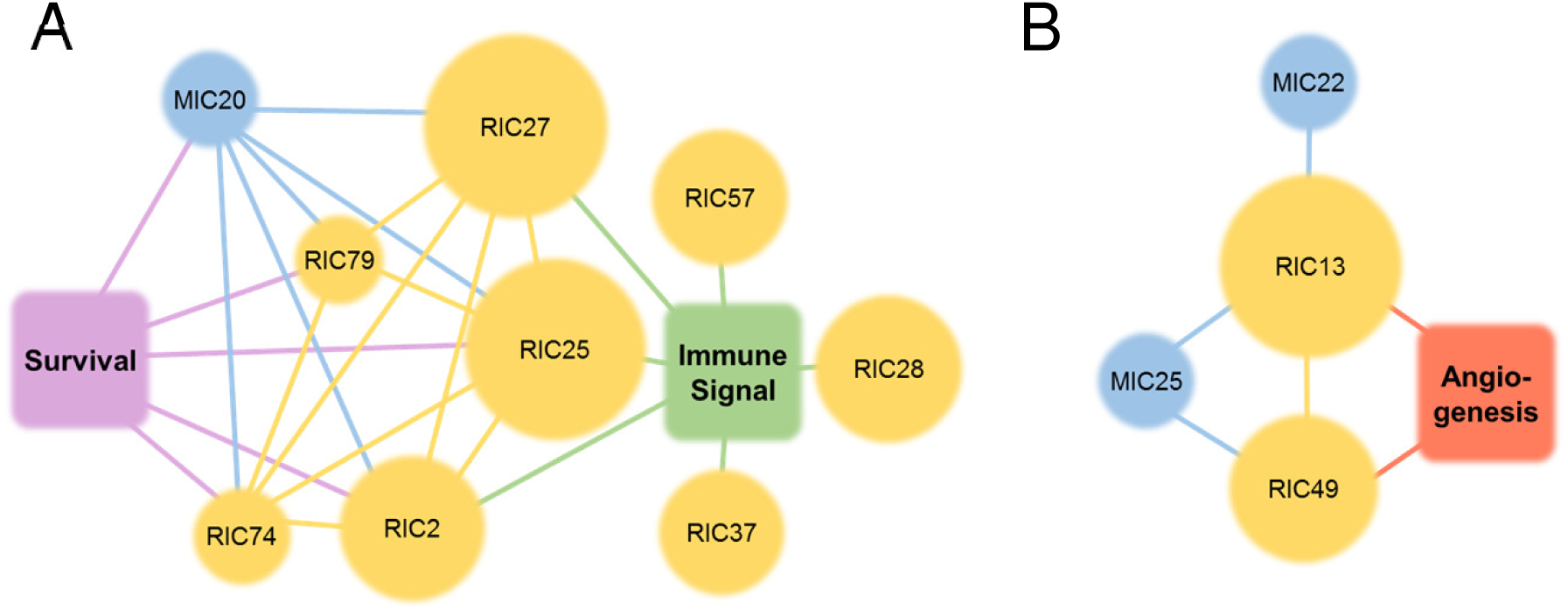
Component clusters. Cluster A is based on gene components (RICs) linked to immune response via enrichment analysis of top-contributing genes; cluster B is based on RICs linked to angiogenesis and stroma transcriptional signal. The size of the circles illustrates the number of top-contributing genes and miRNAs in the components. RIC and MIC components have been linked to each other via correlation analysis (edges between components show absolute correlation over 0.5). Survival analysis was performed by Cox regression with weights of the components used as predictors. See also Supplementary Results (online Supplementary Data) for the details about the components.

Immune component RIC2 was strongly linked to survival (LHR=-0.89, p-value=1.8e-4) and most probably originated from B cells (*Enrichr* ‘B cells’ category enriched, adj.p-value=3.9e-6). The metagenes of RIC2 were also correlated with the LM22 signatures for B cells (‘plasma cells’ r=0.63, ‘B cells memory’ r=0.58, ‘B cells naïve’ r=0.57, Supplementary Results, and also showed the highest correlation with B cell profiles measured in single cells (Supplementary Figure S3)). RIC27 showed a very similar collection of enriched gene sets. These two components most probably represent naïve and activated B-cells.

Functionally, RIC28 was linked to inflammatory responses to wounding (adj.p-value=6.3e-22), neutrophil degranulation (adj.p-value=1.3e-7), TNF (adj.p-value=4.7e-8) and IL1-mediated signalling pathways (adj.p-value=2.2e-9); RIC37 was connected to interferon signalling (adj.p-value = 5.1e-22) whose metagenes were also reciprocally correlated with M5_INTERFERON of the Biton dataset (Biton et al., 2014). Components RIC74 and RIC79 contained a very limited number of top-contributing genes, but both were significantly linked to survival (p-values of 1.3e-3 and 3.2e-3). No specific cell type was associated with these components. RIC74 was, however, associated with positive and negative regulation of immune response and receptor-mediated endocytosis (all adj.p-values=2.6e-4).

The weights of miRNA component MIC20 were positively correlated with the weights of RIC2, RIC25 and RIC27 (correlation of 0.69, 0.86 and 0.64 accordingly) and were positively linked with survival (LHR= −1.32, p-value=1.2e-4). Among the top miRNAs in MIC20 were miR-155, miR-150, miR-342, miR-146b, and miR-142. MiR-155 is known to be a regulator of immune response in cancer cells (Huffaker, Lee et al., 2017, Ji, Wrzesinski et al., 2015) while miR-150, miR-155 and miR-342 have been proposed as markers for melanoma patient survival (Segura, Belitskaya-Levy et al., 2010). Interestingly, four of those positively contributing miRNAs formed a cluster on chr1q32.2 (adj.p-value 7.3e-3).

The new samples were characterised by the involvement of the above immune response-related components (Figure 4A). The results are presented in Table 3. All components linked to subpopulations of immune cells (RIC2, RIC25, RIC57, MIC20) showed little involvement in the new patients suggesting moderate overall immune reactions to the tumour except specific interferon responses, which had high weights in the new samples (RIC28, RIC37).

**Table 3.** Biologically relevant components and their ranked weights in the new samples (investigation dataset). Rank is calculated in comparison to the TCGA reference set (red – weight is above majority of TCGA samples, blue - below). Risk is assigned using Cox regression and log-rank p-value is reported. For MICs the enriched cytogenic bands (adj.p-value<0.05) are presented.

| Cluster | Component | Risk (p-value) | Meaning | P2PM | P4PM | P6PM | P4NS | NHEM |
| --- | --- | --- | --- | --- | --- | --- | --- | --- |
| Immune | RIC2 | decreased (1.8e-4) | B cells | 0.11 | 0.07 | 0.02 | 0.19 | 0.01 |
|  | RIC25 | decreased (2.8e-7) | T cells | 0.26 | 0.06 | 0.24 | 0.18 | 0.00 |
|  | RIC27 | no effect | B cells | 0.80 | 0.37 | 0.31 | 0.80 | 0.00 |
|  | RIC28 | no effect | response to wounding | 0.34 | 0.57 | 0.78 | 0.43 | 0.84 |
|  | RIC37 | no effect | IFN signalling pathway | 0.97 | 0.66 | 0.99 | 0.90 | 1.00 |
|  | RIC57 | no effect | monocytes | 0.00 | 0.25 | 0.24 | 0.02 | 0.00 |
|  | MIC20 | decreased (1.2e-4) | T cells, chr1q32.2 | 0.14 | 0.08 | 0.37 | 0.02 | 0.19 |
| Stromal and angiogenic | RIC13 | no effect | cells of stroma | 0.81 | 0.40 | 0.50 | 0.86 | 0.03 |
|  | RIC49 | no effect | endothelial cells | 0.73 | 0.12 | 0.29 | 0.84 | 0.00 |
|  | MIC22 | no effect | miR-379/miR-410 cluster, chr14q32.2,14q32.31 | 0.29 | 0.20 | 0.27 | 0.38 | 0.16 |
|  | MIC25 | no effect | potentially related to stromal cells; clusters: chr1q24.3, 5q32, 17p13.1, 21q21.1 | 0.97 | 0.85 | 0.76 | 0.80 | 0.26 |
| Skin-related | RIC5 | increased (5.8e-3) | epidermis development and keratinisation | 0.92 | 0.93 | 0.96 | 0.92 | 0.87 |
|  | RIC7 | increased (8.9e-6) | epidermis development and keratinisation | 0.94 | 0.93 | 0.93 | 0.95 | 0.57 |
|  | RIC19 | increased (4.0e-2) | epidermis development and keratinisation | 1.00 | 0.62 | 0.22 | 1.00 | 0.93 |
|  | RIC31 | increased (2.2e-2) | epidermis development and keratinisation | 0.98 | 0.85 | 0.89 | 0.99 | 0.28 |
|  | MIC9 | increased (2.9e-2) | skin-specific miRNAs | 0.95 | 0.88 | 0.87 | 0.91 | 0.83 |
| Melanocytes | RIC4 | increased (5.4e-3) | melanin biosynthesis | 0.62 | 0.77 | 1.00 | 0.21 | 0.96 |
|  | RIC16 | decreased (5.1e-4) | melanosomes (negative gene list) | 0.68 | 0.77 | 0.54 | 0.75 | 0.39 |
|  | MIC11 | no effect | potential regulators of malignant cells, chrXq27.3 | 0.21 | 0.96 | 0.62 | 0.13 | 0.48 |
|  | MIC14 | decreased (1.5e-2) | potential regulators of melanocytes, chrXq26.3 | 0.01 | 0.29 | 0.67 | 0.29 | 0.38 |
| Other | RIC55 | increased (3.0e-2) | cell cycle | 0.48 | 0.46 | 0.88 | 0.00 | 0.53 |
|  | RIC6 | decreased (5.5e-3) | potentially linked to neuron differentiation | 0.43 | 0.73 | 0.59 | 0.46 | 0.01 |
|  | MIC1 | increased (9.4e-4) | regulators of EMT | 0.11 | 0.07 | 0.02 | 0.19 | 0.01 |

#### Stromal and angiogenic components

The second cluster of RICs was linked to the signals of stromal cells and showed enrichment in genes related to angiogenesis. It included four correlated components: RIC13, RIC49, MIC22 and MIC25 (Figure 4B, Supplementary Results). Genes of component RIC13 were enriched in extracellular matrix organisation (adj.p-value = 2e-26) and vasculature development (adj.p-value = 5e-23). The component’s metagenes were strongly correlated with metagene M3_SMOOTH_MUSCLE of Biton et al. (Biton et al., 2014). In the single cell study, the highest correlation of RIC13 metagenes was observed with cancer-associated fibroblasts. Most probably, this component is linked to cells of tumour stroma. Another component from this cluster, RIC49, showed enrichment in GO-terms linked to blood-vessel development and angiogenesis (both with adj.p-value = 6e-24). Its most correlated single cell type was endothelial cells, which also form part of the tumour microenvironment. Thirteen of the positively contributing miRNAs from MIC22 were strongly concentrated in a narrow genomic region in chr14q32.2 (adj.p-value 5.8e-11). MiRNAs of MIC25 were significantly enriched in four cytogenetic locations: chr1q24.3, chr5q32, chr17p13.1 and chr21q21.1 (adj.p-values of 5.0e-6, 2.6e-3, 4.1e-02 and 9.7e-5, respectively).

In the new clinical samples, the highest amount of stromal and endothelial cells were observed in P2PM and P4NS samples. The primary cell line NHEM showed almost no signal of stromal cells. Interestingly, MIC25 was heavily weighted in all new patient samples, excluding this cell line.

#### Skin-related components

RIC5, RIC7, RIC19, RIC31 all showed an enrichment in GO terms related to processes of the skin including epidermis development (adj.p-value < 2e-15 for all mentioned components) and keratinisation (adj.p-value < 1.4e-10). *Enrichr* suggested that the signals of these components are specific to skin (adj.p-value < 1e-50). The dataset contained 48 keratins and many of them were observed among the top-contributing genes: 20 for RIC5, 28 (RIC7), 30 (RIC19) and 13 (RIC31). RIC5 and RIC7 were negatively correlated with survival, which is in concordance with previous observations (Cancer Genome Atlas, 2015). MIC9 with the skin-specific miR-203 (Yi, Poy et al., 2008), was linked to RIC5, RIC7 and RIC31. Furthermore, several components (RIC4, RIC16, MIC11 and MIC14) were connected to the activity of melanocytes. Top-contributing genes of RIC4 were enriched in the melanin biosynthesis process (adj.p-value 1.2e-5) and *Enrichr* linked these genes to melanocytes (adj.p-value 2.8e-25). RIC14 showed an inverse correlation of the weights with RIC4. Both components were linked to survival, but with an opposite effect: while RIC4 increased the risk (LHR=0.18, p-value=5.4e-3), RIC16 increased the survival (LHR= −0.23, p-value=5.1e-4) (Supplementary Results). Many positively contributing miRNAs of the MIC11 component (16 of 33) - a miRNA cluster associated with early relapse in ovarian cancer patients (Bagnoli, De Cecco et al., 2011)-were located on chrXq27.3 (adj.p-value<1e-7).

#### Other tumour-related components

Some components could be linked to transcriptional signals and regulation of cancer cells. For example, RIC55 captured the cell cycle process (adj.p-value 6.6e-29) and the majority of the 383 genes positively associated to this component are known to be involved in cell cycle control with tumour cells contributing the most to cell division activities. Increased cell proliferation was linked to survival (Cox p-value=3.0e-2). In the investigated samples, the highest weight was observed for the most aggressive tumour P6PM and the lowest value for normal skin P4NS.

Several RICs showed linkage to neural tissue. As an example, both positive and negative top-contributing genes of RIC6 were linked to brain in the ARCHS4 tissue sets of *Enrichr* (both adj.p-values < 1e-33). This component was as well associated with patient survival (Cox p-value=5.5e-3). The component indicates the ability of melanoma cells to show expression patterns specific for cells of the neural crest of human embryos and can be linked to motility of malignant melanocytes.

MiRNA component MIC1 showed an interesting bi-modal distribution in the reference dataset (see two clusters in Figure 2D) and was strongly linked to patient survival (Cox p-value=9.4e-4), suggesting two subgroups of melanoma patients with different prognosis. This component most probably was linked to regulation of EMT, as many miRNA positively or negatively influencing the component are known to be EMT regulators or linked to metastasis formation: miR-551, miR-206, miR-34a, miR-1269, miR-205, miR-876, miR-301b, and miR-365a. Based on our analysis of the reference TCGA dataset, these miRNA listed in Supplementary Results can be further investigated as potential survival markers for melanoma patients.

### ICA-derived biological networks

Given the promising results with regard to immune- and angiogenesis-related components, we performed text mining on the terms “B-cell, miRNA and/or cluster”, “T-cell, miRNA and/or cluster” and “angiogenesis, miRNA and/or cluster”, and compiled a list of published miRNAs involved in immune responses and angiogenesis. For the shared top-contributing miRNAs from MIC20, 22, and 25 (Figure 4 and Supplementary Results), experimentally confirmed target genes were extracted (from miRTarBase (Chou, Shrestha et al., 2018)). In order to investigate possible miRNA-target gene interactions as an underlying biological reasoning for clustering, we next overlaid the extracted target genes with gene lists of connected RICs (Figure 4). Finally, enrichment analysis was performed and final gene lists were analysed by STRING (Szklarczyk, Morris et al., 2017) to visualise potential protein-protein interactions. Overall, all networks (Figure 5, Supplementary Figures S4 and S5) showed a significant enrichment of interactions suggesting a non-random relation between top-contributing miRNAs and genes. STRING network analysis captured key biological interactions reflecting the ICA-based RICs and MICs, from which they were initially derived.

**Figure 5.**
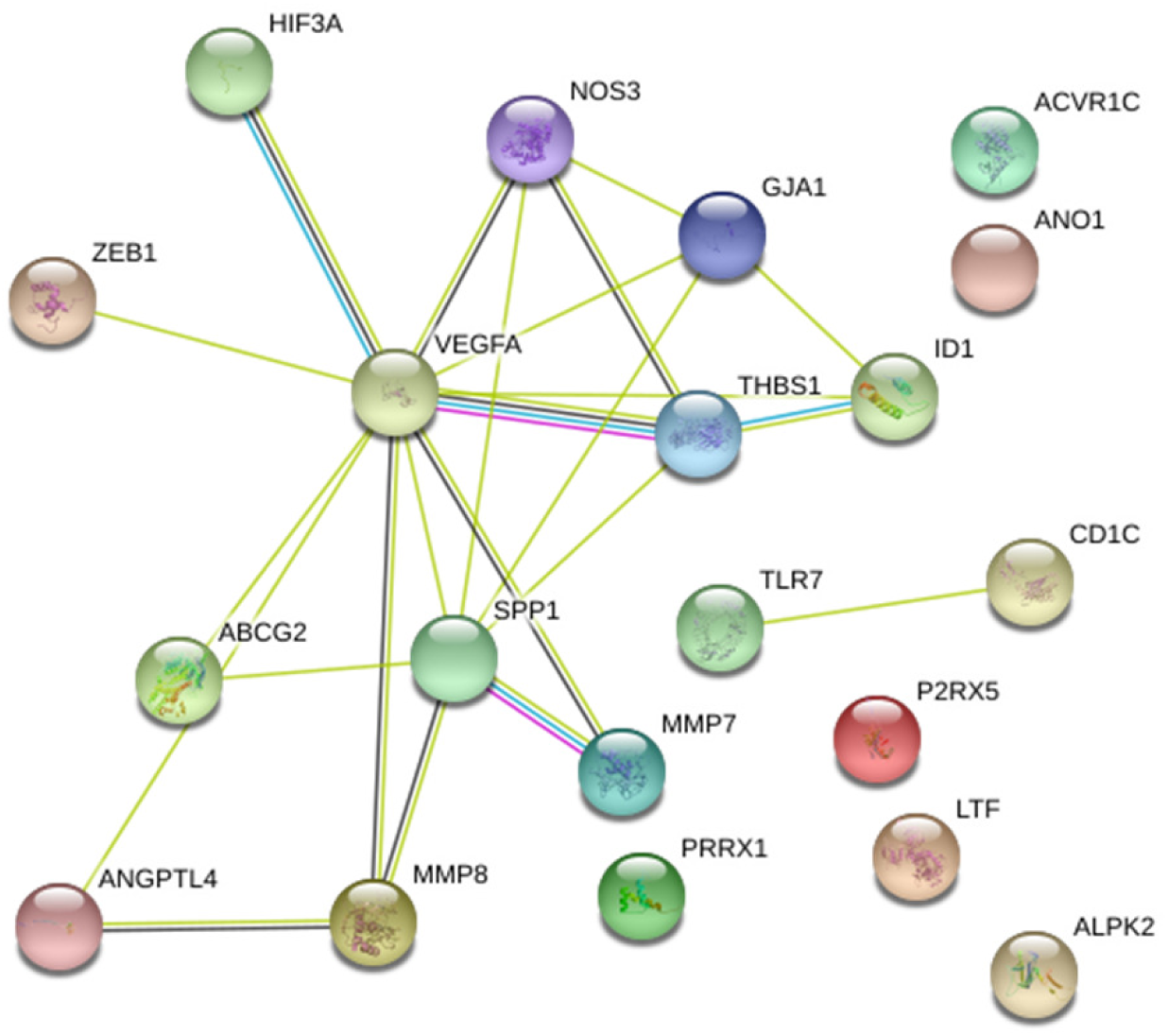
STRING network based on overlapping MIC22-target genes and RIC13 metagenes, showing a significant protein interaction network (PPI enrichment p-value: 1.27e-13) representing main players within angiogenesis. The gene list uploaded into STRING represents the overlap between the target genes of top-contributing miRNAs of MIC22 found in angiogenesis-related publications and top-contributing genes of RIC13 (also see Figure S2, red overlap in the green box for biological relevance).

## DISCUSSION

Here we investigated the applicability of ICA-based deconvolution of transcriptomes, originated from a large set of bulk melanoma samples, for acquiring clinically and biologically relevant information about new patients. ICA decomposes transcriptomics data into components that are characterised by two matrices: a matrix of metagenes, which shows how each gene is involved in each component, and the matrix of weights that represents the involvement of the components in each sample. Importantly, this analysis does not require any preliminary knowledge about biology or sample composition. Unlike other deconvolution methods that use signatures (Yoshihara, Shahmoradgoli et al., 2013) or pure transcriptomic profiles (Quon, Haider et al., 2013), ICA is an assumption-free, unsupervised approach. The method directly works with the data from bulk samples without any preliminary assumption about the transcriptomes of the purified cell types. Among the components, one can expect to see not only those defined by ‘pure’ tumours or stromal cells, but also those originating from tumour/stroma interactions including tumour-induced stromal cell reprogramming. One example of such interactions is angiogenesis, further discussed below.

We implemented a robust consensus ICA method and applied it to several datasets from patients with cutaneous melanoma (SKCM). These included (a) a large cohort of SKCM patients from TCGA, used as reference set; (b) an independent cohort of 44 patients with publicly available microarray mRNA data and (c) 5 in-house clinical investigation samples: 3 primary melanomas, a normal skin sample and a normal melanocyte cell line (NHEM). Both mRNA and miRNA datasets were obtained for the reference and investigation samples. Despite the fact that different techniques were used for data acquisition, ICA was able to identify common signals in the datasets and properly allocate the new samples within the reference set (Figure 2). This was particularly evident for miRNA data where the reference set was obtained by small RNA-seq and the new samples by qPCR arrays with PCA showing a strong difference between these two datasets. With ICA, technical biases in the data were isolated within several components and thus separated from biologically relevant signals leading to a better and more correct characterisation of the samples.

The fact that ICA should be rerun for every series of new samples could be considered as a drawback of our approach. However, recalculation of the components does not require supervision and could be done automatically. In the case when investigation and reference datasets come from the same distribution, one can use the matrix ***S*** obtained from the reference dataset in order to define the weights (***M***) for the samples forming the investigation dataset (Eq.1). However, in reality, the variability in the data requires recalculation of the components for the new investigated samples.

We demonstrate here that when analysing data from melanoma patients, the weights of independent components can be used as predictive features of patient subgroups and can also be linked to patient survival. While the ICA-based feature extraction method has been previously discussed (e.g. (Aziz, Verma et al., 2016, Teschendorff et al., 2007)), no studies have been devoted, to our knowledge, to estimating patient prognosis using ICA-based data deconvolution. We combined weights of several significant components into a hazard score, for which a high predictive power was shown both in the reference cohort (460 patients with known survival status) and in the independent validation cohort (44 patients). Thus, the developed approach could help clinicians in estimating the risks and potentially optimising the selection of adequate treatment strategies. Three of the survival-associated components were connected to immune response. As expected, higher immune signal indicated lower risk for the patients (Bogunovic et al., 2009). Interestingly, two of the skin-related components were as well linked to survival; however, their presence increased the risk, which is in agreement with previous observations (Cancer Genome Atlas, 2015).

Next, the biological relevance of the components was examined in depth. Components that represented signals from various cell subpopulations (e.g. different immune cells, stromal cells, melanocytes) and cellular processes (e.g. cell cycle) were identified. These signals were also detected in the new samples, providing hints of active processes and tissue composition of these samples. We associated mRNA and miRNA components that showed similar weight profiles in all the patients and hypothesised that such components were probably derived from the same cell types or process. For example, MIC20 was correlated with RIC2 and RIC25 – the components associated with leukocyte activity. Indeed, miR-155, one of the markers of immune cells (Emming, Chirichella et al., 2018), was found among the most involved miRNAs of MIC20. Therefore, we could link all other top-contributing miRNAs within MIC20 to leukocytes and immune response and thus assign functions to these miRNAs.

Another group of components were assigned to tumour-stromal interactions and angiogenesis. One of them, MIC22, contained an almost complete miRNA mega cluster miR-379/miR-410, with 11 of 13 miRNAs significantly involved. The cluster is located on chromosome 14 (14q32) in the so-called imprinted DLK1-DIO3 region. Lower levels of this miRNA cluster have been described to favour neo-vascularisation (Welten, Bastiaansen et al., 2014) and shown to play a role in development, neonatal metabolic adaption but also in tumorigenesis. Deregulation of miRNAs in this locus has recently been shown to predict lung cancer patient outcome (Enfield, Martinez et al., 2016, Valdmanis, Roy-Chaudhuri et al., 2015). Most miRNAs in this cluster (68%) were significantly downregulated in glioblastoma multiform, 61% downregulated in kidney renal clear cell carcinoma and 46% in breast invasive carcinoma indicating a tumour suppressive role of this miRNA cluster, especially in glioblastomas (Laddha, Nayak et al., 2013). Moreover, Zehavi et al. (Zehavi, Avraham et al., 2012) have shown that the miR-379/miR-410 cluster was silenced in melanoma, which favoured tumorigenesis and metastasis.

Overall, we observed that ICA on miRNA expression data grouped together many miRNAs that belong to genetic clusters and by connecting MICs with genes (RICs), biological functions of miRNAs could be inferred. As an example, MIC11 represents a cluster on chrX q27.3 associated with early relapse in advanced stage ovarian cancer patients (Bagnoli et al., 2011). In our analysis, the miRNAs from that cluster were linked to activity of malignant melanocytes. All this is suggestive of a concerted role for miRNAs of a given cluster in regulating functionally related genes (Haier, Strose et al., 2016, Wang, Luo et al., 2016).

The results for the ICA-derived biological networks implied that the combination of ICA with text mining (biological expressions enriched in statistically correlated RICs and MICs) potentially uncovers two hidden connections: biological reasons for statistical correlations and detection of those genes actually responsible for the biological link between MICs and RICs. This in turn might give new insights into the significance of biological processes involved in cancer in general or in certain cancer subtypes.

Similarly to PCA, ICA could be integrated into standard analysis pipelines in the future. Unlike PCA, which only groups samples in the space of a few principle components, ICA could extract biologically-based signals. These signals can be further used to acquire clinically relevant information about new samples, thus helping patient diagnostics and prognostics. Taken together, consensus ICA approach represents a versatile tool to dissect complex data cohorts into individual components allowing for better use of such datasets.

## MATERIAL AND METHODS

### Preparation of the reference and validation datasets

#### Expression data

As reference dataset, we used the open-access TCGA skin cutaneous melanoma (SKCM) datasets, namely RNA-seq (HTSeq raw counts, FPKM and TPM) and miRNA-seq data (miRNA isoform read count) from the Genomic Data Commons (GDC) data portal of the National Cancer Institute of the National Institutes of Health (NIH, https://portal.gdc.cancer.gov/). The RNA-seq dataset comprises data from 468 different individuals (472 samples). Of those, 368 originated from metastatic samples (1 individual provided 2 samples) and 103 from primary melanoma tumours; one sample represented a solid normal tissue. MiRNA-seq data were available for 452 individuals, with 353 metastatic, 97 primary tumour and 2 normal skin tissue samples. The miRNA-isoform read counts data were collapsed per isoform and IDs were mapped to miRNA-names based on miRBase v. 21 (http://www.mirbase.org/).

A validation dataset of gene expression data was taken from Bogunovic et al. (Bogunovic et al., 2009), available from ArrayExpress under E-GEOD-19234. This Affymetrix GeneChip Human Genome U133 Plus 2.0 microarray dataset consisted of 44 metastatic samples from melanoma patients accompanied by survival information. As microarray expression data have very different dynamic range compared to RNA-seq (Nazarov, Muller et al., 2017), we shifted and scaled the microarray data: the 5^th^ percentile of expression was used as the lowest meaningful signal and was subtracted from microarray gene expression. All negative values were set to 0. The data were then scaled to unify the 75^th^ percentile between reference RNA-seq and validation microarray data.

#### Clinical data

To explore the possibility of assigning clinical traits to TCGA samples, we compiled a small dataset based on public TCGA data with “fail-safe” items covering gender and sample type (primary tumour and metastatic). Additionally we added publication-based data for RNA-seq clustering (immune / keratin / MITF-low) (Cancer Genome Atlas, 2015) as this information has been claimed to be relevant for disease prognosis. Survival data were extracted by parsing related information out of publicly available individual clinical data files (XML files provided by GDC). We extracted and processed the information assigned to the tags: bcr_patient_barcode, days_to_death, vital_status, year_of_initial_pathologic_diagnosis, age_at_initial_pathologic_diagnosis, days_to_last_followup and person_neoplasm_cancer_status. The full survival and clinical datasets are described in Supplementary Tables S3 and S4, respectively.

### Preparation of the investigation dataset: clinical samples, data acquisition and analysis

The investigation dataset, represented by RNA-seq and miRNA qPCR array data, is composed of primary tumour samples of three melanoma patients and two control samples (one matched normal skin and a healthy melanocyte cell line, NHEM). Sample annotation is presented in Table 1. Details of sample collection, preparation, transcriptome and miRNome analyses are described in Supplementary Methods. RNA-seq data for these samples are available by GEO accession number GSE116111 and Ct-values for all quantified miRNAs are available in Supplementary Table S5.

To harmonise miRNA annotation of qPCR arrays and TCGA-derived miRNA isoform read count data, we first re-annotated our qPCR arrays to miRNA version 21. To have comparable data between qPCR arrays and TCGA, we worked with miRNA isoform data referring to miRNA IDs, so that mapping of stem loop IDs to mature miRNA IDs was possible.

### Data Analysis

#### RNA-seq expression measures

Four metrics of gene expression were considered: raw counts, DESeq2-normalized counts (Anders & Huber, 2010), Fragments Per Kilobase of transcript per Million (FPKM) and Transcripts Per kilobase Million (TPM). All expression values were log_2_ transformed. Raw gene expression data were represented by 60446 features, of which many were lowly expressed. In order to reduce the number of uninformative features, we applied soft filtering, by the maximum expression level as described in (Nazarov, Reinsbach et al., 2013): only genes that showed over 1000 counts in at least one sample of the reference TCGA SKCM dataset were considered (Supplementary Figure S6A shows distribution of maximum gene expression and the threshold in log_2_ scale). This resulted in 16579 informative genes (distribution is presented in Supplementary Figure S6B). ICA of raw counts showed the best performance for patient stratification with smaller number of components using gender and sample type as benchmark (described in Supplementary Materials, Figure S6C and S6D).

#### Independent component analysis

Independent component analysis (ICA) was applied to the combined reference and investigation datasets for unsupervised separation of signals and feature extraction (Figure 1). By combining the datasets, we expect that technical biases between the reference and investigation data are estimated by the method and isolated within some of the components. Each layer of omics data: mRNA and miRNA was analysed independently at this stage. ICA implementation from the *fastICA* package of R was used (Marchini, Heaton et al., 2017). Let us denote ***E*_*nm*_** the expression matrix of *n* genes or miRNAs measured in *m* bulk samples. ICA decomposed such a matrix into a product of *k* statistically independent transcriptional signals ***S*_*nk*_** (addressed further as matrix of metagenes) and a weight or mixing matrix ***M*_*km*_** (matrix of metasamples) (Hyvarinen, 1999, Kairov, Cantini et al., 2017, Zinovyev, Kairov et al., 2013).

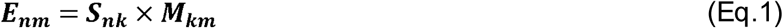

The values represented in the columns of ***S*** (metagenes) can be interpreted as the level of influence of the corresponding genes/miRNAs on the components. When a component captures a specific cell type, its corresponding set of top-contributing genes can be considered as markers of this cell type. Weights in rows of ***M*** show how the metagenes are mixed in the samples. Again, if a component is linked to a cell type, its weight may be considered as an estimation of the fraction of those cells in each sample. In order to distinguish independent components obtained after ICA of mRNA and miRNA data, we introduce the terms RICs (mRNA) and MICs (miRNAs). Thus, each RIC and MIC is associated with two vectors: one shows the involvement of the genes in this component (a column of ***S***); the second represents the weights of the component in the samples (a row of ***M***). Unlike non-negative matrix factorization, both metagenes and weights can be positive or negative and *ab initio* the selection of the direction is random, depending on the initial estimation. Because of this, ICA may suffer from reduced reproducibility for at least some components. To mitigate this drawback, we ran the analysis multiple times (100 runs during the exploratory step and 1000 for the final analysis). The detailed algorithm of consensus ICA is described in the Supplementary Methods. Multithreading was implemented in R code to speed-up calculations using the *foreach* package and either *doMC* (Linux) or *doSNOW* (MS Windows) packages available in R/Bioconductor. The script of the implemented consensus ICA and other tools used here for investigation of the components are available online: https://gitlab.com/biomodlih/consica.

The top-contributing genes and miRNAs per component were detected using the following significance analysis approach. A p-value was independently assigned to each gene/miRNA within each component, based on the probability that it came from a normal distribution with estimated parameters. As a small subset of genes had extremely high values in ***S***, while the majority was normally distributed, we used non-parametric measures to estimate the centre and scale of the distribution (median and median absolute deviation). Then these p-values were corrected for multiple testing (Benjamini & Hochberg), and those with an adj.p-value<0.01 were reported as top-contributing genes of a component. Two lists of top-contributing genes/miRNAs resulted from the analysis – positively and negatively involved. The lists of top-contributing genes of each RIC were afterwards used for over-representation (enrichment) analysis. The 16579 informative genes were used as a background gene list and significantly enriched (adj.p-value<0.01) GO terms were investigated. In order to simplify the interpretation and to increase the robustness for runs on different datasets, we reoriented the components in order to have the most significantly enriched categories associated with positive top-contributing genes (Supplementary Methods). For MICs, the direction could not be identified by enrichment analysis, therefore we reoriented only those MICs that showed strong negative correlation with RICs. As we ran ICA on a combination of reference and investigation datasets, several components captured the platform difference between these datasets. We labelled such components as technical components. Accordingly, the remaining components could be considered as being clean from this confounding effect.

#### Prediction of new sample classes

Random forest (RF) classifier, implemented in the *randomForest* R package (Liaw & Wiener, 2002), was used with the default settings to predict classes of patients. Columns of the weight matrix ***M*** were used as inputs and clinical variables (e.g. gender, sample type) as outputs. Each variable was analysed independently. First, leave-one-out cross-validation (LOOCV) was performed on the reference set in order to address the ability of predicting sample classes and estimate the accuracy of prediction. Then the RF, trained on all reference data, was used to predict classes for the new clinical samples of the investigation dataset.

#### Integration of mRNA and miRNA expression data for survival prediction

Weights of the components (rows of matrix ***M***) were statistically linked to patient survival using Cox partial hazard regression implemented in the *survival* package of R (Therneau & Grambsch, 2000). FDR-adjusted p-values of the log rank test were used to select significant components. However, the prognostic power of each individual component might not have been high enough to be applied to the patients from the new cohort. Therefore, we integrated weights of several components, calculating the hazard score (HS) with an improved predictive power. For each patient, its HS is the sum of the products of log hazard ratio (LHR) of the univariable Cox regression (LHR, only significant), the component stability R^2^ (see Supplementary Methods) and the standardised row of weight matrix ***M***:

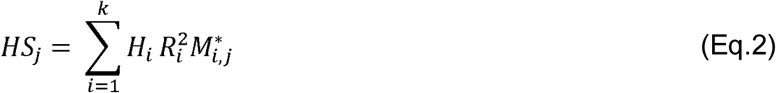

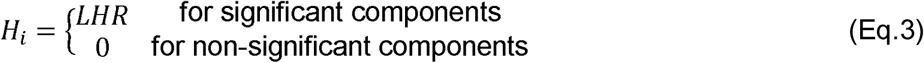

The validity of the proposed score was checked using the validation set.

#### Biological relevance of the components

Our strategy to investigate the biological relevance of the components is presented in Supplementary Figure S1. First, we attempted to connect the metagenes of all the components from the mRNA data to biological functions and cell types. We analysed separately the positively and negatively involved genes using several tools. Automatic analysis was done by *topGO* (Alexa & Rahnenfuhrer, 2016) followed by a manual analysis with *Enrichr* (Kuleshov et al., 2016, http://amp.pharm.mssm.edu/Enrichr/) that checked for enrichment in multiple categories originated from various databases (we used Reactome 2016, GO Biological Processes 2017, Human Gene Atlas, ARCHS4 Tissues and Chromosome Location, http://amp.pharm.mssm.edu/Enrichr/#stats). In addition, we compared the metagenes to the ones previously published by Biton et al. (Biton et al., 2014) and assigned the component number to the reciprocally corresponding metagene as explained in Cantini et al. (Cantini, Kairov et al., 2018) using the *DeconICA* R package (https://zenodo.org/record/1250070). As enrichment of immune-related processes and functions was observed, we also correlated our metagenes to the immune cell type signature matrix named *LM22* (Newman et al., 2015) in order to identify components originated from different types of leukocytes; cell-types were associated with components through highest correlation. Finally, for some components we confirmed their biological origin by correlating the metagenes with averaged gene expression profiles of cell types measured at a single-cell level and reported by Tirosh et al. (Tirosh et al., 2016). For miRNA data we considered enrichment of genomic locations of contributing miRNAs annotated by the *cyto_convert* tool of NCBI (https://www.ncbi.nlm.nih.gov/genome/tools/cyto_convert/). Standard hypergeometric tests with p-value adjustment was used to associate over-representation of miRNAs from MICs within cytogenic bands.

#### Integration of components for data at miRNA and mRNA levels

Pearson correlation between weights of the components was used to link the components found within mRNA and miRNA data. Here we hypothesized that if two components show significant correlation of the weights in all the samples, they should be functionally linked. Of note, these MICs have been linked to their respective RIC, purely based on the high correlation of component weights, without considering any biological knowledge. Due to the lack of tools providing data with regard to biological functions or cell types for miRNAs, we performed literature mining, searching for all publications related to miRNAs-clusters and additional biologically relevant keywords. Based on intermediate results suggesting a connection of network-related RICs to specific cell types, namely T- and B-cells or angiogenesis, we used these expressions as keywords, assuming a biological link between MICs and RICs. After automatically extracting miRNA-names and clusters from publication titles and abstracts by an inhouse Python script (available on demand), we compared those with the miRNA-metagenes comprised in the correlation-based networks, to identify a possible enrichment of miRNAs related to the proposed biological function. Additionally, we compared our miRNA-metagenes to *miRCancer* (http://mircancer.ecu.edu/ (Xie, Ding et al., 2013)).

In order to explore the link or edges between MICs and RICs we extracted the target genes with a strong support from *miRTarBase* (http://mirtarbase.mbc.nctu.edu.tw/php/index.php, (Chou et al., 2018)) for those miRNA-metagenes mapping the miRNAs and clusters found by literature mining. Additionally, we filtered these target genes to ensure that they were part of the reference gene set based on top-contributing genes as determined by the mRNA-reference set. We then overlapped these target genes with the metagenes of the respective linked RIC and applied *Enrichr* tool through the automated Python-based API using the following reference gene set collections: KEGG_2016, GO_Biological_Process_2017b, GO_Cellular_Component_2017b, Jensen_TISSUES, Jensen_DISEASES (script available on demand). Moreover, we explored the overlapping target- and metagenes by STRING (https://string-db.org/, (Szklarczyk et al., 2017)) to detect significantly enriched protein-protein interaction networks. Both the results of the link-analysis between RICs and MICs as well as the investigation of MIC-metagenes have been finalised by visual inspection and evaluation by biological experts. A detailed schematic illustration is shown in Supplementary Figure S1.

The involvement or the weight of each component in the new samples was not centred and scaled. To visualize the involvement of the components in the new samples, we replaced the weights of the components by a ranking score that changed from 0 to 1 (only reference set data were considered to define the ranking). If the weight of the considered component in a new sample was below (or above) the weights in the reference set, such component automatically was assigned to a limiting value of 0 (or 1). Values of ranking score around 0.5 in the new sample suggest that the weight of the considered component was close to the median in the reference set.

## DATA AVAILABILITY

### RNA-seq and miRNA-seq of investigation set

GDC data portal: https://portal.gdc.cancer.gov/

### Expression data of validation set

Array Express: https://www.ebi.ac.uk/arrayexpress/experiments/E-GEOD-19234/

### New expression data of investigation set

The sequencing data for 3 primary melanoma tumours and 2 controls are freely available under the GEO accession number GSE116111. Data for miRNAs are in the Supplementary Table S5. *Tools:*

Consensus parallel ICA: https://gitlab.com/biomodlih/consica

R/Bioconductor v.3.4.3 with packages *fastICA, doMC, doSNOW, topGO, randomForest*,

*survival* https://cran.r-project.org/

*Enrichr* http://amp.pharm.mssm.edu/Enrichr/

*STRING* https://string-db.org/

*miRTarBase* http://mirtarbase.mbc.nctu.edu.tw/php/index.php

*miRBase* http://www.mirbase.org/

*miRCancer* http://mircancer.ecu.edu/

*DeconICA* https://github.com/sysbio-curie/DeconICA

cyto_convert https://www.ncbi.nlm.nih.gov/genome/tools/cyto_convert/

## SUPPLEMENTARY DATA

Supplementary materials that include Supplementary Figures, Supplementary Tables, Supplementary Methods and Supplementary Results are available online.

## ACKNOWLEDGEMENT

We would like to thank the patients for providing clinical material and the clinical staff of the Dermatology Unit at the University of Freiburg for professionally taking and handling all the samples. Bioinformatics analyses presented in this paper were partly carried out using the high-performance computing facilities of the University of Luxembourg (http://hpc.uni.lu). Furthermore, we thank Demetra Philippidou and Dr. Susanne Reinsbach for processing clinical samples and NHEM cells forming the investigation dataset. We acknowledge Dr. Aurélien Ginolhac and Cristina Maximo for critically reading the manuscript and for valuable input.

## REFERENCES

Alexa A, Rahnenfuhrer J (2016) topGO: Enrichment analysis for Gene Ontology. In

Anders S, Huber W (2010) Differential expression analysis for sequence count data. Genome biology 11: R106

Aziz R, Verma CK, Srivastava N (2016) A fuzzy based feature selection from independent component subspace for machine learning classification of microarray data. Genom Data 8: 4–15

Bagnoli M, De Cecco L, Granata A, Nicoletti R, Marchesi E, Alberti P, Valeri B, Libra M, Barbareschi M, Raspagliesi F, Mezzanzanica D, Canevari S (2011) Identification of a chrXq27.3 microRNA cluster associated with early relapse in advanced stage ovarian cancer patients. Oncotarget 2: 1265–78

Biton A, Bernard-Pierrot I, Lou Y, Krucker C, Chapeaublanc E, Rubio-Perez C, Lopez-Bigas N, Kamoun A, Neuzillet Y, Gestraud P, Grieco L, Rebouissou S, de Reynies A, Benhamou S, Lebret T, Southgate J, Barillot E, Allory Y, Zinovyev A, Radvanyi F (2014) Independent component analysis uncovers the landscape of the bladder tumor transcriptome and reveals insights into luminal and basal subtypes. Cell reports 9: 1235–45

Bogunovic D, O’Neill DW, Belitskaya-Levy I, Vacic V, Yu YL, Adams S, Darvishian F, Berman R, Shapiro R, Pavlick AC, Lonardi S, Zavadil J, Osman I, Bhardwaj N (2009) Immune profile and mitotic index of metastatic melanoma lesions enhance clinical staging in predicting patient survival. Proceedings of the National Academy of Sciences of the United States of America 106: 20429–34

Cancer Genome Atlas N (2015) Genomic Classification of Cutaneous Melanoma. Cell 161: 1681–96

Cantini L, Kairov U, De Reynies A, Barillot E, Radvanyi F, Zinovyev A (2018) Stabilized Independent Component Analysis outperforms other methods in finding reproducible signals in tumoral transcriptomes. BioRxiv

Chou CH, Shrestha S, Yang CD, Chang NW, Lin YL, Liao KW, Huang WC, Sun TH, Tu SJ, Lee WH, Chiew MY, Tai CS, Wei TY, Tsai TR, Huang HT, Wang CY, Wu HY, Ho SY, Chen PR, Chuang CH et al. (2018) miRTarBase update 2018: a resource for experimentally validated microRNA-target interactions. Nucleic Acids Res 46: D296–D302

Debey S, Schoenbeck U, Hellmich M, Gathof BS, Pillai R, Zander T, Schultze JL (2004) Comparison of different isolation techniques prior gene expression profiling of blood derived cells: impact on physiological responses, on overall expression and the role of different cell types. The pharmacogenomics journal 4: 193–207

Emming S, Chirichella M, Monticelli S (2018) MicroRNAs as modulators of T cell functions in cancer. Cancer Lett 430: 172–178

Enfield KS, Martinez VD, Marshall EA, Stewart GL, Kung SH, Enterina JR, Lam WL (2016) Deregulation of small non-coding RNAs at the DLK1-DIO3 imprinted locus predicts lung cancer patient outcome. Oncotarget 7: 80957–80966

Haier J, Strose A, Matuszcak C, Hummel R (2016) miR clusters target cellular functional complexes by defining their degree of regulatory freedom. Cancer Metastasis Rev 35: 289-322

Hayward NK, Wilmott JS, Waddell N, Johansson PA, Field MA, Nones K, Patch AM, Kakavand H, Alexandrov LB, Burke H, Jakrot V, Kazakoff S, Holmes O, Leonard C, Sabarinathan R, Mularoni L, Wood S, Xu Q, Waddell N, Tembe V et al. (2017) Whole-genome landscapes of major melanoma subtypes. Nature 545: 175–180

Holderfield M, Deuker MM, McCormick F, McMahon M (2014) Targeting RAF kinases for cancer therapy: BRAF-mutated melanoma and beyond. Nat Rev Cancer 14: 455–67

Huffaker TB, Lee SH, Tang WW, Wallace JA, Alexander M, Runtsch MC, Larsen DK, Thompson J, Ramstead AG, Voth WP, Hu R, Round JL, Williams MA, O’Connell RM (2017) Antitumor immunity is defective in T cell-specific microRNA-155-deficient mice and is rescued by immune checkpoint blockade. J Biol Chem 292: 18530–18541

Hyvarinen A (1999) Fast and robust fixed-point algorithms for independent component analysis. IEEE Trans Neural Netw 10: 626–34

Ji Y, Wrzesinski C, Yu Z, Hu J, Gautam S, Hawk NV, Telford WG, Palmer DC, Franco Z, Sukumar M, Roychoudhuri R, Clever D, Klebanoff CA, Surh CD, Waldmann TA, Restifo NP, Gattinoni L (2015) miR-155 augments CD8+ T-cell antitumor activity in lymphoreplete hosts by enhancing responsiveness to homeostatic gammac cytokines. Proceedings of the National Academy of Sciences of the United States of America 112: 476–81

Kairov U, Cantini L, Greco A, Molkenov A, Czerwinska U, Barillot E, Zinovyev A (2017) Determining the optimal number of independent components for reproducible transcriptomic data analysis. BMC Genomics 8: 712

Kuleshov MV, Jones MR, Rouillard AD, Fernandez NF, Duan Q, Wang Z, Koplev S, Jenkins SL, Jagodnik KM, Lachmann A, McDermott MG, Monteiro CD, Gundersen GW, Ma’ayan A (2016) Enrichr: a comprehensive gene set enrichment analysis web server 2016 update. Nucleic Acids Res 44: W90–7

Laddha SV, Nayak S, Paul D, Reddy R, Sharma C, Jha P, Hariharan M, Agrawal A, Chowdhury S, Sarkar C, Mukhopadhyay A (2013) Genome-wide analysis reveals downregulation of miR-379/miR-656 cluster in human cancers. Biol Direct 8: 10

Lawrence MS, Stojanov P, Polak P, Kryukov GV, Cibulskis K, Sivachenko A, Carter SL, Stewart C, Mermel CH, Roberts SA, Kiezun A, Hammerman PS, McKenna A, Drier Y, Zou L, Ramos AH, Pugh TJ, Stransky N, Helman E, Kim J et al. (2013) Mutational heterogeneity in cancer and the search for new cancer-associated genes. Nature 499: 214–218

Lee SI, Batzoglou S (2003) Application of independent component analysis to microarrays. Genome biology 4: R76

Legres LG, Janin A, Masselon C, Bertheau P (2014) Beyond laser microdissection technology: follow the yellow brick road for cancer research. American journal of cancer research 4: 1–28

Liaw A, Wiener M (2002) Classification and regression by randomForest. R News 2: 18–22

Luke JJ, Flaherty KT, Ribas A, Long GV (2017) Targeted agents and immunotherapies: optimizing outcomes in melanoma. Nat Rev Clin Oncol 14: 463–482

Marchini JL, Heaton C, Ripley BD (2017) fastICA: FastICA Algorithms to Perform ICA and Projection Pursuit. In

Nazarov PV, Muller A, Kaoma T, Nicot N, Maximo C, Birembaut P, Tran NL, Dittmar G, Vallar L (2017) RNA sequencing and transcriptome arrays analyses show opposing results for alternative splicing in patient derived samples. BMC Genomics 18: 443

Nazarov PV, Reinsbach SE, Muller A, Nicot N, Philippidou D, Vallar L, Kreis S (2013) Interplay of microRNAs, transcription factors and target genes: linking dynamic expression changes to function. Nucleic Acids Res 41: 2817–31

Newman AM, Liu CL, Green MR, Gentles AJ, Feng W, Xu Y, Hoang CD, Diehn M, Alizadeh AA (2015) Robust enumeration of cell subsets from tissue expression profiles. Nat Methods 12: 453–7

Patel AP, Tirosh I, Trombetta JJ, Shalek AK, Gillespie SM, Wakimoto H, Cahill DP, Nahed BV, Curry WT, Martuza RL, Louis DN, Rozenblatt-Rosen O, Suva ML, Regev A, Bernstein BE (2014) Single-cell RNA-seq highlights intratumoral heterogeneity in primary glioblastoma. Science 344: 1396–401

Quon G, Haider S, Deshwar AG, Cui A, Boutros PC, Morris Q (2013) Computational purification of individual tumor gene expression profiles leads to significant improvements in prognostic prediction. Genome Med 5: 29

Schadendorf D, Fisher DE, Garbe C, Gershenwald JE, Grob JJ, Halpern A, Herlyn M, Marchetti MA, McArthur G, Ribas A, Roesch A, Hauschild A (2015) Melanoma. Nat Rev Dis Primers 1: 15003

Segura MF, Belitskaya-Levy I, Rose AE, Zakrzewski J, Gaziel A, Hanniford D, Darvishian F, Berman RS, Shapiro RL, Pavlick AC, Osman I, Hernando E (2010) Melanoma MicroRNA signature predicts post-recurrence survival. Clin Cancer Res 16: 1577–86

Shannon CP, Balshaw R, Ng RT, Wilson-McManus JE, Keown P, McMaster R, McManus BM, Landsberg D, Isbel NM, Knoll G, Tebbutt SJ (2014) Two-stage, in silico deconvolution of the lymphocyte compartment of the peripheral whole blood transcriptome in the context of acute kidney allograft rejection. PloS one 9: e95224

Sullivan RJ, Flaherty KT (2013) Resistance to BRAF-targeted therapy in melanoma. Eur J Cancer 49: 1297–304

Szklarczyk D, Morris JH, Cook H, Kuhn M, Wyder S, Simonovic M, Santos A, Doncheva NT, Roth A, Bork P, Jensen LJ, von Mering C (2017) The STRING database in 2017: quality-controlled protein-protein association networks, made broadly accessible. Nucleic Acids Res 45: D362–D368

Taroni JN, Greene CS (2017) Cross-Platform Normalization Enables Machine Learning Model Training On Microarray And RNA-Seq Data Simultaneously. bioRxiv

Teschendorff AE, Journee M, Absil PA, Sepulchre R, Caldas C (2007) Elucidating the altered transcriptional programs in breast cancer using independent component analysis. PLoS computational biology 3: e161

Therneau TM, Grambsch PM (2000) Modeling survival data: extending the Cox model. Springer, New York

Tirosh I, Izar B, Prakadan SM, Wadsworth MH, 2nd, Treacy D, Trombetta JJ, Rotem A, Rodman C, Lian C, Murphy G, Fallahi-Sichani M, Dutton-Regester K, Lin JR, Cohen O, Shah P, Lu D, Genshaft AS, Hughes TK, Ziegler CG, Kazer SW et al. (2016) Dissecting the multicellular ecosystem of metastatic melanoma by single-cell RNA-seq. Science 352: 189–96

Valdmanis PN, Roy-Chaudhuri B, Kim HK, Sayles LC, Zheng Y, Chuang CH, Caswell DR, Chu K, Zhang Y, Winslow MM, Sweet-Cordero EA, Kay MA (2015) Upregulation of the microRNA cluster at the Dlk1-Dio3 locus in lung adenocarcinoma. Oncogene 34: 94–103

Wang Y, Luo J, Zhang H, Lu J (2016) microRNAs in the Same Clusters Evolve to Coordinately Regulate Functionally Related Genes. Mol Biol Evol 33: 2232–47

Weaver WM, Tseng P, Kunze A, Masaeli M, Chung AJ, Dudani JS, Kittur H, Kulkarni RP, Di Carlo D (2014) Advances in high-throughput single-cell microtechnologies. Current opinion in biotechnology 25: 114–23

Welten SM, Bastiaansen AJ, de Jong RC, de Vries MR, Peters EA, Boonstra MC, Sheikh SP, La Monica N, Kandimalla ER, Quax PH, Nossent AY (2014) Inhibition of 14q32 MicroRNAs miR-329, miR-487b, miR-494, and miR-495 increases neovascularization and blood flow recovery after ischemia. Circ Res 115: 696–708

Xie B, Ding Q, Han H, Wu D (2013) miRCancer: a microRNA-cancer association database constructed by text mining on literature. Bioinformatics 29: 638–44

Yi R, Poy MN, Stoffel M, Fuchs E (2008) A skin microRNA promotes differentiation by repressing ‘stemness’. Nature 452: 225–9

Yoshihara K, Shahmoradgoli M, Martinez E, Vegesna R, Kim H, Torres-Garcia W, Trevino V, Shen H, Laird PW, Levine DA, Carter SL, Getz G, Stemke-Hale K, Mills GB, Verhaak RG (2013) Inferring tumour purity and stromal and immune cell admixture from expression data. Nat Commun 4: 2612

Zehavi L, Avraham R, Barzilai A, Bar-Ilan D, Navon R, Sidi Y, Avni D, Leibowitz-Amit R (2012) Silencing of a large microRNA cluster on human chromosome 14q32 in melanoma: biological effects of mir-376a and mir-376c on insulin growth factor 1 receptor. Mol Cancer 11: 44

Zhang T, Dutton-Regester K, Brown KM, Hayward NK (2016) The genomic landscape of cutaneous melanoma. Pigment Cell Melanoma Res 29: 266–83

Zhao B, Hemann MT, Lauffenburger DA (2014) Intratumor heterogeneity alters most effective drugs in designed combinations. Proceedings of the National Academy of Sciences of the United States of America 111: 10773–8

Zinovyev A, Kairov U, Karpenyuk T, Ramanculov E (2013) Blind source separation methods for deconvolution of complex signals in cancer biology. Biochemical and biophysical research communications 430: 1182–7

